# FtsZ of wall-less bacteria forms ring-like structures

**DOI:** 10.1101/2024.10.15.618391

**Authors:** Taishi Kasai, Yohei O Tahara, Makoto Miyata, Daisuke Shiomi

**Affiliations:** Department of Life Science, College of Science, Rikkyo University; Graduate School of Science, Osaka Metropolitan University; The OMU Advanced Research Center for Natural Science and Technology, Osaka Metropolitan University

**Keywords:** cell division, FtsZ, SepF, *Spiroplasma*, L-form

## Abstract

The FtsZ protein is involved in bacterial cell division. In cell-walled bacteria, such as *Bacillus subtilis*, FtsZ forms a ring-like structure, called the Z ring, at the cell division site and acts as a scaffold for cell wall synthesis. The inhibition of cell wall synthesis in *B. subtilis* has been shown to interfere with the function of the Z ring, causing a loss in cell division control. *Spiroplasma*, a cell wall-less bacterium, lacks most of the genes involved in cell division; however, the *ftsZ* gene remains conserved. The function of *Spiroplasma eriocheiris* FtsZ (SeFtsZ) remains to be determined. In the present study, we analyzed the biochemical characteristics of SeFtsZ. Purified SeFtsZ demonstrated lower polymerization capacity and GTPase activity than FtsZ from *E. coli* and *B. subtilis*. We also investigated the relationship between SeFtsZ and SeSepF, which anchors FtsZ to the cell membrane, and found that SeSepF did not contribute to the stability of FtsZ filaments, unlike the *B. subtilis* SepF. SeFtsZ and SeSepF were produced in *E. coli* L-forms, where cell wall synthesis was inhibited. SeFtsZ formed ring-like structures in cell wall-less *E. coli* cells, suggesting that SeFtsZ forms Z rings and is involved in cell division independently of cell wall synthesis.

## Introduction

Bacteria generally divide at the middle of the cell. The cell division apparatus complex called the Z ring, of which FtsZ is a major component, must be correctly localized at the division site. FtsZ is a tubulin homolog consisting of a globular tubulin-like domain containing GTP-binding and GTPase domains, an H7 domain that connects GTP-binding and GTPase domains, a T7 loop, an intrinsically disordered C-terminal “linker,” and a C-terminal tail which is a highly conserved ∼11 residues region responsible for the interaction with other proteins. The T7 loop from one FtsZ subunit is inserted into the GTP-binding site of the next, inducing GTP hydrolysis (1). The tense state (T-state) of FtsZ bound to GTP polymerizes to form filaments. FtsZ hydrolyzes GTP in the filament, changing it to the relaxed state (R-state); in turn, the filament becomes destabilized and depolymerized (2). Leucine residue at the 272 position in the FtsZ globular domain in *Escherichia coli* is homologous to residue involved in the longitudinal contact of β-tubulin (3, 4). When this residue is mutated to glutamate (L272E), FtsZ fails to polymerize and loses its GTPase activity (5, 6).

In *Bacillus subtilis*, the Z ring localization at mid-cell is determined by nucleoid occlusion and the Min system (7, 8). Noc is a protein involved in nucleoid occlusion (9, 10). It is thought to recruit chromosomes to the cell membrane and physically prevent the assembly of the cell division machinery near the nucleoid (11). In contrast, MinC prevents the assembly of the cell division machinery at the cell poles (12). The Z ring is anchored to the cell membrane by FtsA and SepF, which interact with the C-terminus of FtsZ and stabilize its filament (13, 14). The actin homolog FtsA has an amphipathic helix at its C-terminus, a membrane-bound region, and SepF is self-assembled and forms a ring structure (15–18). The SepF ring bundles FtsZ filaments and transforms them into tube-like structures (16). Glycine residues at positions 109 and 116 of SepF in *B. subtilis* are important for interacting with FtsZ (16). Gly109 also affects intermolecular interactions with SepF dimers (16). Once anchored to the cell membrane, the Z-ring assembles with other cell division proteins to form the complex ‘divisome,’ which acts as a scaffold for cell wall synthesis (19, 20). FtsW and PBP2B function as a transglycosylase and transpeptidase, respectively (21, 22). FtsL, DivIB, and DivIC form complexes involved in the localization of FtsW and PBP2B to the Z ring (23).

In cell-walled bacteria, such as *B. subtilis*, cell wall synthesis is linked to cell division since the dividing site must be filled with a new cell wall (24, 25). Cell wall synthesis inhibition has been shown to interfere with the function of cell division proteins. For example, when cell wall synthesis is inhibited by β-lactam antibiotics, such as penicillin, *E. coli* and *B. subtilis* cells are lysed since β-lactams inhibit the activity of penicillin-binding proteins (PBPs), which crosslinks the peptides in peptidoglycan (26). However, under hyperosmotic conditions, although most cells are lysed owing to the inhibition of cell wall synthesis by antibiotics or lysozymes, some cells are transformed into a viable state without a cell wall, referred to as the L-form. In the L-forms, FtsZ is not required for cell division, and the elongated cell membranes divide irregularly (27). However, even in *E. coli* L-forms, the Z ring is formed but does not constrict, indicating that peptidoglycan or its synthesis is required for the constriction of the Z ring (Hayashi et al., in revision). Consistently, *E. coli* FtsZ inside liposomes forms Z-rings that do not split liposomes (28). Therefore, FtsZ-regulated cell division in cell-walled bacteria may require the cell wall. However, some bacteria, such as *Mycoplasmas*, are not originally surrounded by a cell wall and, as in the case of *Mycoplasma genitalium,* encode the *ftsZ* gene in their genomes, although *Mycoplasma* lacks most of the genes involved in cell wall synthesis (29, 30). *ftsZ* deficiency in *Mycoplasma* affects cell morphology and division rates (31). Synthetic bacterium JCVI-Syn3.0, which has the smallest genome derived from *Mycoplasma*, showed an increased growth rate when several genes, including FtsZ, were introduced (32). Therefore, *Mycoplasma* FtsZ may function differently from the FtsZ of *E. coli* and *B. subtilis*. However, data regarding cell division in bacteria without cell walls, other than in cell-walled bacteria such as *E. coli* or *B. subtilis*, and the regulation of FtsZ localization and its functions during cell division remain limited.

The function of cytoskeletal proteins is altered in *Spiroplasma*, a close relative of *Mycoplasma*. MreB, a scaffold for cell-wall synthesis during cell elongation in *E. coli* and *B. subtilis*, is the internal structure that maintains the cell morphology of *Spiroplasma* (33, 34). In this study, we investigated the role of FtsZ in *Spiroplasma* cell division and, using purified FtsZ of *Spiroplasma eriocheiris* (SeFtsZ), determined the biochemical properties and structure of SeFtsZ. We also used walled or wall-deficient (L-form) *E. coli* cells to observe the behavior of SeFtsZ in cells.

## Results

### GTP-dependent polymerization of *Spiroplasma* FtsZ

His_6_-SeFtsZ and His_6_-SeSepF (hereafter referred to as SeFtsZ and SeSepF, respectively, unless otherwise indicated) were overproduced in *E. coli* and purified using His-tag affinity chromatography. To analyze the polymerization of SeFtsZ, purified SeFtsZ was incubated in a polymerization buffer with GTP or GDP for 30 min. Samples were fractionated via ultracentrifugation. The amount of SeFtsZ in the supernatant and pellet fractions was estimated from the bands of Coomassie brilliant blue (CBB)-stained SDS-PAGE gel (Fig. 1). Almost all SeFtsZ was detected in the supernatant in the presence of GDP, whereas 16% of SeFtsZ was retrieved from the pellet in the presence of GTP, consistent with previous findings (35, 36). Next, the effect of SeSepF on SeFtsZ polymerization was examined. Purified SeFtsZ was mixed with purified SeSepF in polymerization buffer for 5 min. The mixed solution was then incubated with GTP or GDP for 30 min. Almost all SeFtsZ and SeSepF were detected in the supernatant in the presence of GDP. In the presence of GTP, 28% of SeFtsZ was retrieved from the pellet fraction, suggesting that SeSepF promotes the polymerization of SeFtsZ.

**Figure 1.**
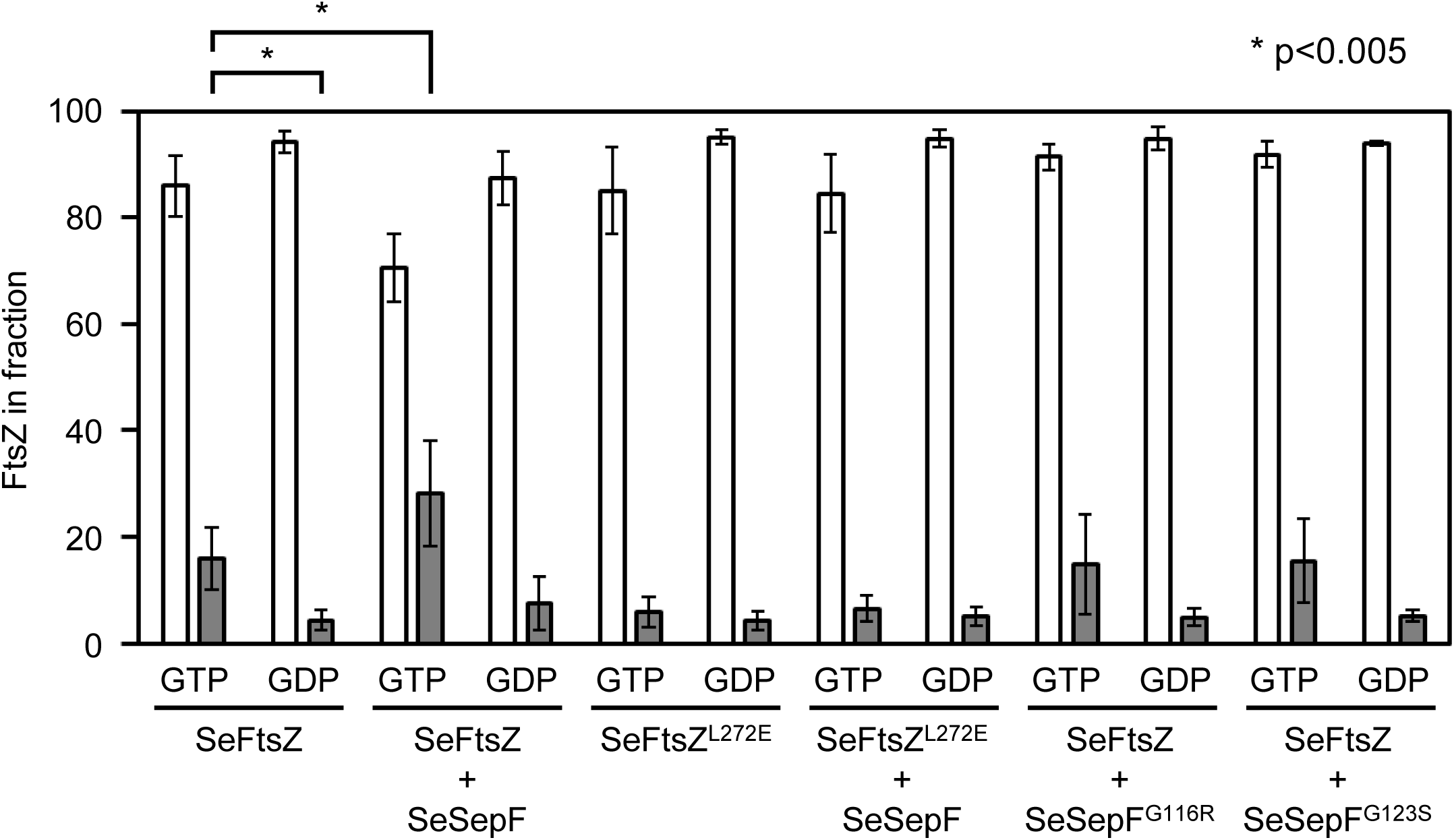
Sedimentation assay of SeFtsZ polymer. SeFtsZ (12 μM) was polymerized with 2 mM GTP or GDP. The amount of protein was estimated using densitometric analysis of CBB-stained SDS-PAGE gels (12%).

To confirm that the increase in SeFtsZ sedimentation was attributed to the interaction between SeFtsZ and SeSepF, we first examined the interaction between SeFtsZ and SeSepF using bio-layer interferometry. Before the analysis, the His-tag of SeFtsZ was removed since one of the His-tagged protein (His_6_-SeSepF) had to be immobilized on a chip. His-tagged SeSepF was then immobilized on a Ni-NTA biosensor chip. The dissociation constant (Kd) of SeFtsZ with SeSepF in the absence and presence of GTP were 1.04 and 53.9 μM, respectively (Fig. 2A). Next, we measured the interaction between the SeFtsZ and SeSepF mutants. L272 of FtsZ and G109 and G116 of SepF in *B. subtilis* correspond to L272 of SeFtsZ and G116 and G123 of SeSepF (16). When L272 of SeFtsZ was replaced with E, the SeFtsZ^L272E^ mutant was expected to fail to polymerize and lose GTPase activity. When G116 and G123 of SeSepF were replaced with R and S, respectively, the SeSepF mutants (SepF^G116R^ and SepF^G123S^, respectively) were expected to show decreased interaction with SeFtsZ. The binding affinity of SeFtsZ^L272E^ with SeSepF was not affected in the absence or presence of GTP (Kd = 0.98 μM and 0.7 μM) (Fig. 2B). The concentration-dependent signal of SeFtsZ did not increase in the sensor immobilized with SeSepF^G116R^ and SeSepF^G123S^ mutants (Fig. 2C and 2D), indicating that the interaction between SeFtsZ and SeSepF mutants was lost due to the mutations. The dissociation constant of interaction between SeFtsZ and SeSepF was 1.13 μM as that of *Mycobacterium tuberculosis* (37).

**Figure 2.**
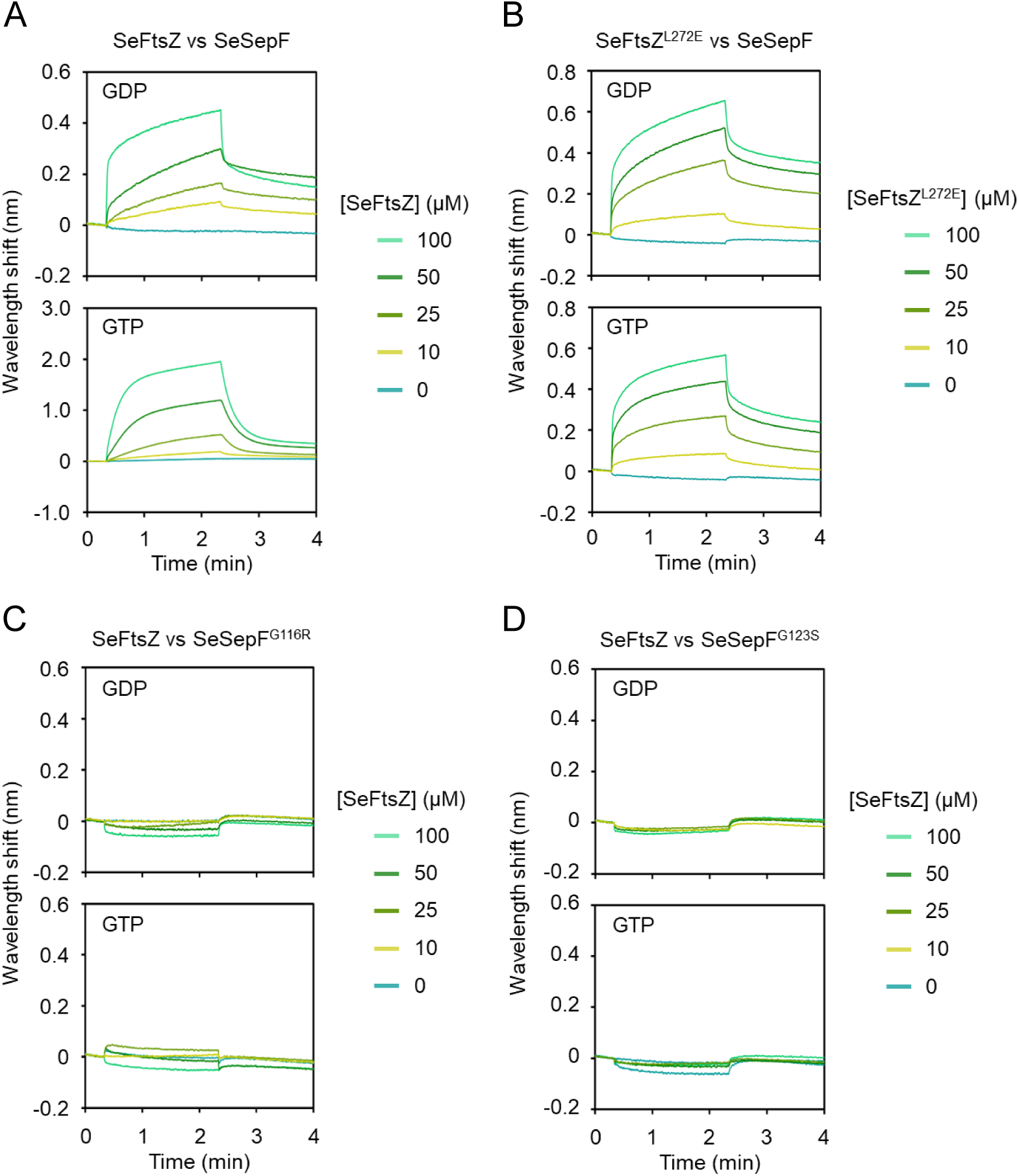
Interaction between SeFtsZ and SeSepF. Binding analysis of SeFtsZ and SeFtsZ^L272E^ mutant with SeSepF and SeSepF mutants was conducted using Bio-layer interferometry. SeSepF (A and B) and SeSepF^G116R^ (C), SeSepF^G123S^ (D) immobilized on the sensor chip were ligands. The concentrations of SeFtsZ (A, C, D) and SeFtsZ^L272E^ (B) mutant, which are the analytes.

SeFtsZ^L272E^ did not polymerize under any of the conditions (Fig. 1). SeSepF mutants (SeSepF^G116R^ and SeSepF^G123S^) were tested for polymerization. The amount of SeFtsZ in the pellet fraction did not increase, regardless of the presence of GTP or SeSepF (Fig. 1). These results indicate that SeSepF promotes SeFtsZ polymerization through direct interactions.

### Observation of *Spiroplasma* FtsZ filaments

Next, we observed the structures formed by polymerized SeFtsZ. Polymerized SeFtsZ was imaged using an electron microscopy (EM). The structures composed of SeFtsZ were thin, short filaments (Fig. 3A). Some filaments are curved into partial circles. The width of the filament was approximately 5.7 ± 1.2 nm, with an inner diameter of the circular structure of 50–100 nm. SeFtsZ was expected to be larger, as determined by EM staining. The SeFtsZ polymer incubated with SeSepF formed thin, short filaments, and some long, thick bundles (Fig. 3B). The SeFtsZ^L272E^ mutant did not form filaments or bundles in the presence or absence of SeSepF (Fig. 3C and 3D). Purified SepF from *B. subtilis* forms tubular structures under neutral pH conditions and ring structures under basic pH (16, 18). We determined whether SeSepF formed any characteristic structures. The structures of SeSepFs were imaged using EM. No SeSepF rings or tubules were observed at neutral pH, suggesting that SeFtsZ filaments are more unstable than *B. subtilis* FtsZ filaments.

**Figure 3.**
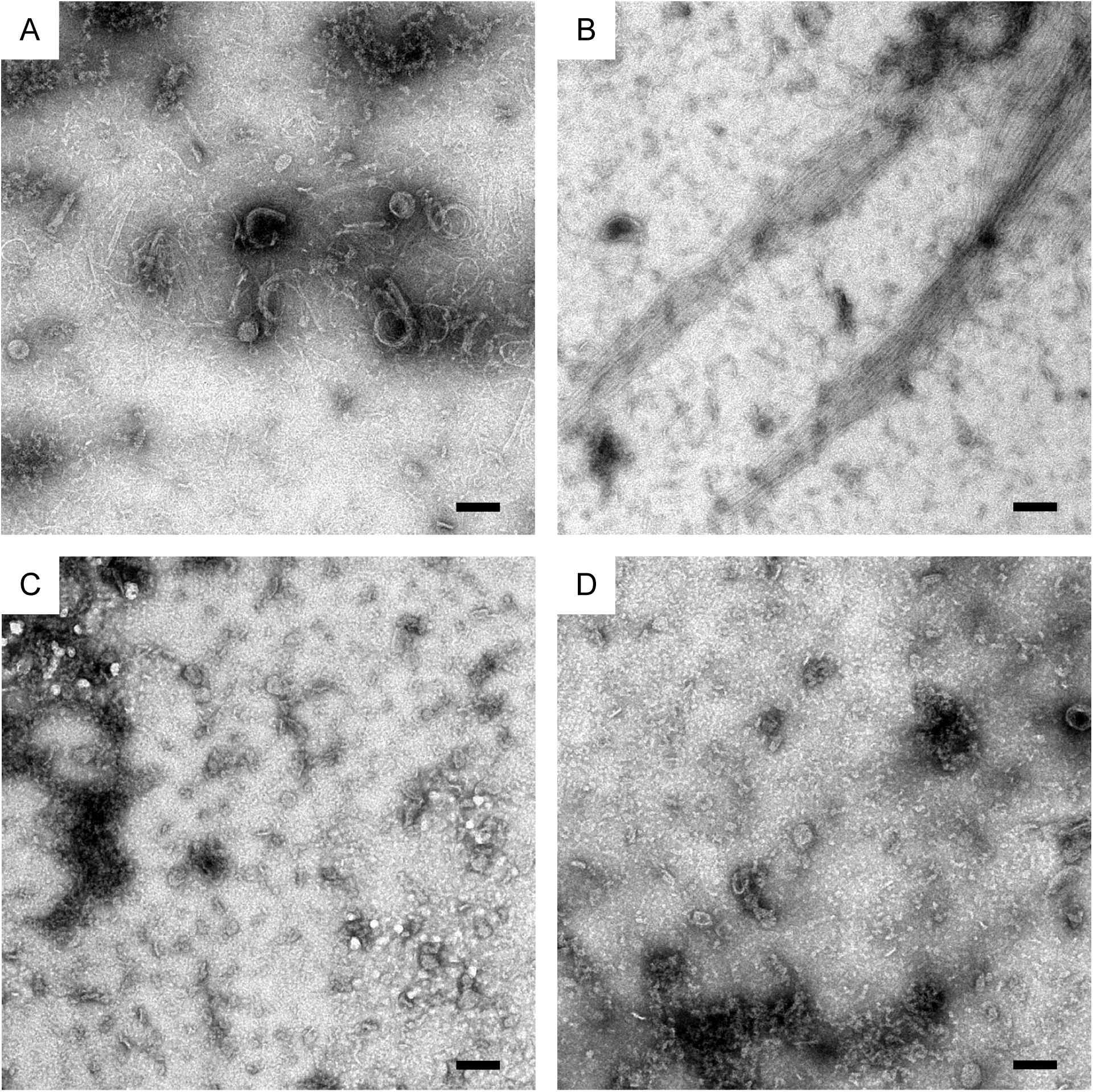
Electron microscope image of polymerized SeFtsZ. SeFtsZ filaments in the absence (A) and presence (B) of SeSepF. SeFtsZ^L272E^ was not polymerized without (C) or with (D) SeSepF. Scale bar: 100 nm.

### The GTPase activity of *Spiroplasma* FtsZ

Next, we analyzed the effects of SeSepF on the GTPase activity of SeFtsZ. SeFtsZ hydrolysis was determined based on the amount of inorganic phosphate. GTP (1 mM) was added to 12 μM SeFtsZ, and the absorbance (OD_620_) of the solution was measured for 60 min. The amount of inorganic phosphate in the reaction solution increased for 20 min after adding GTP. The GTPase activity of SeFtsZ was 0.2 ± 0.04 Pi/FtsZ/min (Fig. 4). The GTPase activity of SeFtsZ was measured in the presence of SeSepF. SeSepF slightly increased the GTPase activity of SeFtsZ to 0.26 ± 0.06 Pi/FtsZ/min (Fig. 4). The SeFtsZ^L272E^ mutant did not exhibit an increase in the amount of inorganic phosphate in the reaction solution after 60 min (Fig. 4). SeSepF increased the GTPase activity of SeFtsZ since polymerized SeFtsZ was increased by SeSepF. The GTP hydrolysis of SeFtsZ was lower than *E. coli* and *B. subtilis* FtsZ. In *B. subtilis*, SepF stabilizes FtsZ filaments and decreases GTPase activity (14). These results suggest that SeSepF does not contribute to the stability of SeFtsZ filaments.

**Figure 4.**
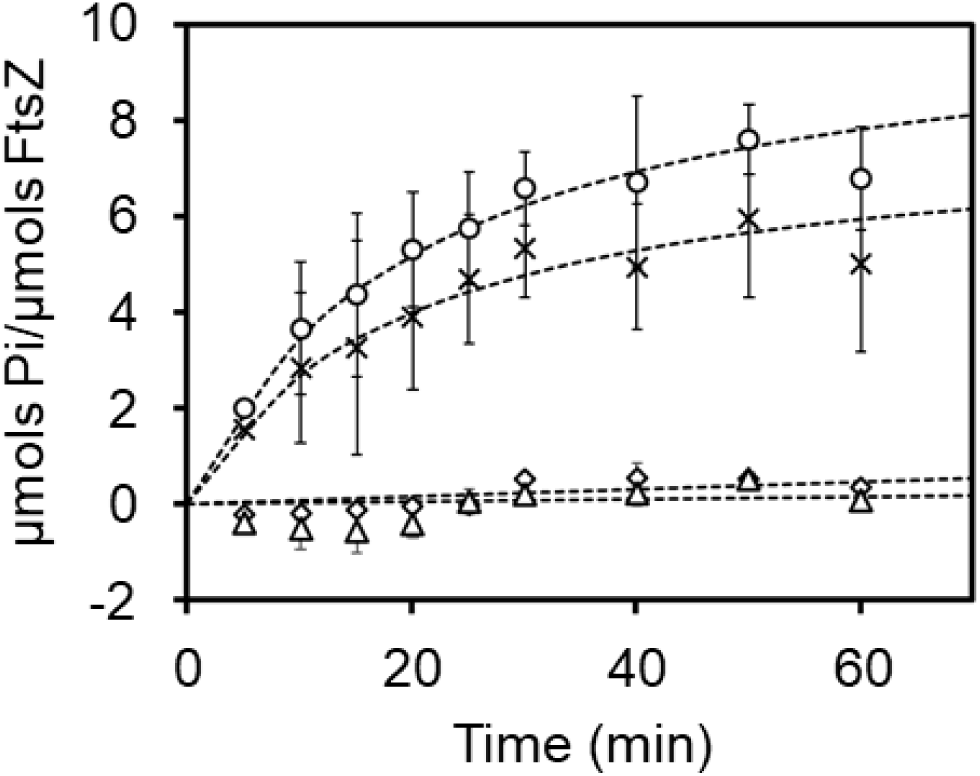
GTP hydrolysis during SeFtsZ polymerization. SeFtsZ (12 μM) polymerized with 2 mM GTP. The reactions in the absence of SeSepF are indicated as cross signs, and those in the presence of SeSepF as circle signs. SeFtsZ^L272E^ (12 μM) incubated in the absence (square) or presence (triangle) with 2 mM GTP.

### Localization and structure of *Spiroplasma* FtsZ in *E. coli*

In general, FtsZ polymers form the Z ring at cell division sites. However, the ring-like structure of SeFtsZ in *S. eriocheiris* cells has not been reported. Herein, SeFtsZ formed a Z-ring in living cells; however, a genetic manipulation technique has not been established for *Spiroplasma*. Since *S. eriocheiris* is an obligate anaerobic bacterium, its cultivation is difficult; even if cultured, its growth rate is very slow. Therefore, it is difficult to observe the Z ring in *S. eriocheiris* cells. Previously, we successfully produced FtsZ derived from *Arabidopsis* chloroplasts in *E. coli* and observed ring-like structures, indicating that *E. coli* cells can form Z-rings derived from other species (38). Hence, in this study, we observed SeFtsZ localization in *E. coli* (Fig. 5). Since FtsZ interacts with other cell division proteins through its C-terminal region, this region may be critical for SeFtsZ function. Therefore, the monomeric superfolder GFP (msfGFP) was fused to the N-terminus of SeFtsZ. When msfGFP-SeFtsZ was expressed in *E. coli*, it formed foci instead of ring-like structures (Fig. 5B). Notably, cells carrying an empty vector did not elongate (Fig. 5A), whereas cells producing SeFtsZ became filamentous (Fig. 5B). These results suggest that SeFtsZ do not assemble into the Z ring of *E. coli* but inhibit cell division possibly by interacting with endogenous cell division proteins in *E. coli* and/or inhibiting a turn-over of EcFtsZ. Next, msfGFP-SeFtsZ was coproduced with SeSepF in *E. coli* cells. msfGFP-SeFtsZ formed ring-like structures and msfGFP-SeFtsZ was excluded from the cell poles (Fig. 5C). The msfGFP-SeFtsZ^L272E^ mutant not only formed foci but also inhibited cell division (Fig. 5D).

**Figure 5.**
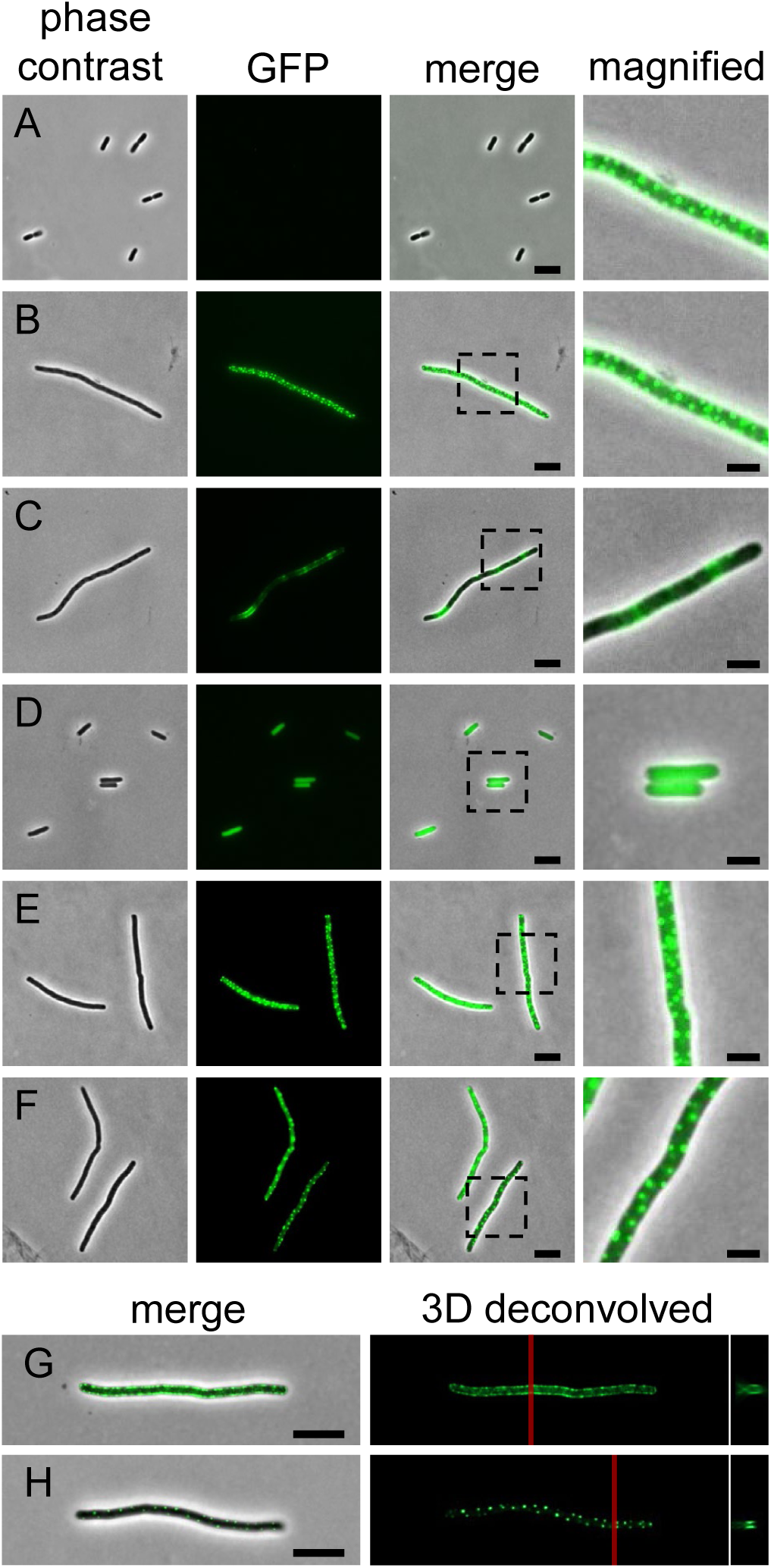
Localization of SeFtsZ in Rod-shaped *E. coli*. (A) MG1655 with introduced vector. (B) The msfGFP-SeFtsZ focus formation in MG1655. (C) The msfGFP-SeFtsZ formed a ring structure in MG1655 expressing SeSepF. (D) The msfGFP-SeFtsZ^L272E^ diffused in the cytoplasm in the presence of SeSepF. (E, F) The msfGFP-SeFtsZ formed a focused structure in MG1655 expressing SeSepF mutant. The magnified cell is shown in the right panel. (G, H) The three-dimensional deconvolved images of msfGFP-SeFtsZ focus and ring. A cross-section of a red line is shown. Scale bar: 2 μm.

SeSepF^G116R^ and SeSepF^G123S^ mutants did not affect the localization of msfGFP-SeFtsZ foci (Fig. 5E and 5F). To confirm whether msfGFP-SeFtsZ forms ring-like structures in a SepF-dependent manner, we analyzed the localization of SeFtsZ in three dimensions, and only msfGFP-SeFtsZ clusters were observed (Fig. 5G); in the presence of SepF, ring-like structures of msfGFP-SeFtsZ were observed (Fig. 5H). Collectively, SeFtsZ polymerized and formed cluster-like foci within living *E. coli* cells and possibly interacted with SeSepF to form ring-like structures. The inhibition of cell division in *E. coli* is thought to be caused by SeFtsZ polymerization.

### Localization of *Spiroplasma* FtsZ in *E. coli* L-forms

msfGFP-SeFtsZ formed ring-like structures in rod-shaped cells. However, the production of msfGFP-SeFtsZ and SeSepF inhibited cell division in cell-walled *E. coli*. The cell wall may affect the contraction of rings containing msfGFP-SeFtsZ. Therefore, we analyzed the localization of msfGFP-SeFtsZ in cell wall-deficient L-form cells and the division of L-forms that produce msfGFP-SeFtsZ (Fig. 6A). When msfGFP-SeFtsZ was produced in the L-form *E. coli*, foci were observed in the cytoplasm. The msfGFP-FtsZ focal point was eventually polymerized and extended. The growth rate of SeFtsZ filaments was 14 ± 6 nm/min (N = 10). SeFtsZ filaments interacted with each other and formed long filaments or branches. Ring- and tube-like SeFtsZ structures were also observed. The average diameter of the msfGFP-SeFtsZ ring had an outer diameter of 500 ± 75 nm and an inner diameter of 215 ± 57 nm. These structures were unlikely to anchor to the cell membrane and did not control the division of L-form cells. The *E. coli* FtsZ ring has a diameter of 1.0 μm (39). The widths of *S. eriocheiris* and *E. coli* are roughly 200 and 1 µm, respectively, suggesting that the curvature of FtsZ filaments depended on the bacterial cell size (40). Next, msfGFP-SeFtsZ and SeSepF were co-produced in L-form cells (Fig. 6B). msfGFP-SeFtsZ localized to the cell membrane and formed a ring-like structure. The msfGFP-SeFtsZ ring appeared to contract the cell membrane in the narrow part of the cell (Fig. 6C), regardless of whether SeFtsZ generated forces to contract the cell or coincidentally localized where the cells thinned out.

**Figure 6.**
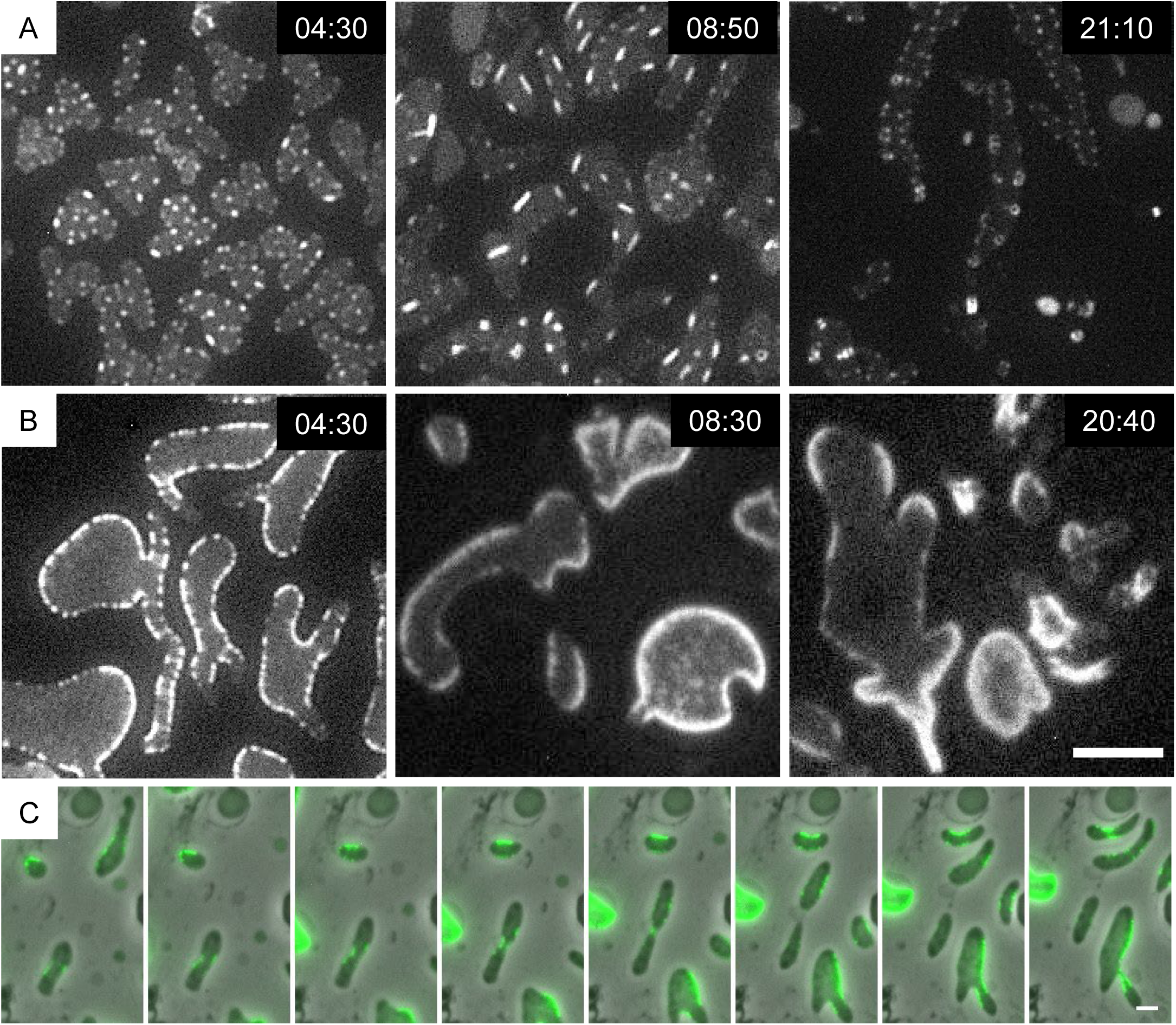
Localization of SeFtsZ in L-form *E. coli*. Time lapse image showing the msfGFP-SeFtsZ behavior. Cells were grown in NB/MSM medium containing 400 μg/mL Fosfomycin and 0.2% arabinose. (A) The msfGFP-SeFtsZ filaments elongated and formed the rings in MG1655. (B) The msfGFP-SeFtsZ localized at the cell membrane in MG1655, expressing SeSepF. Scale bar: 5 μm. (C) Time lapse images of cell division in L-form. L-form divided at the localized portion of the msfGFP-SeFtsZ. Scale bar: 2 μm.

## Discussion

Most studies on bacterial cell division have been conducted on cell-walled bacteria, such as *E. coli* and *B. subtilis*. However, there have been limited studies on cell division in bacteria without cell walls (30–32). The *ftsZ* gene is widely conserved in almost all bacteria, with or without cell walls. Therefore, we predicted that FtsZ is important for cell division irrespective of the presence or absence of a cell wall. However, the mechanism behind FtsZ’s involvement in cell division in bacteria lacking a cell wall and the biochemical characteristics of FtsZ in these bacteria are still unknown. Herein, we report the biochemical characteristics of FtsZ in *Spiroplasma*.

### The GTPase activity of FtsZ

The GTPase activity of *E. coli* and *B. subtilis* FtsZ are 2.1 Pi/FtsZ/min and 0.8 Pi/FtsZ/min, respectively (35). The GTPase activity of SeFtsZ was 0.2 ± 0.04 Pi/FtsZ/min (Fig. 4). Although SeFtsZ contains conserved amino acids essential for GTP binding and hydrolysis, its GTPase activity is lower than that of *E. coli* and *B. subtilis* FtsZ, which is possibly attributed to the phenylalanine residue at position 226 in SeFtsZ. This residue is predicted to affect conformational changes in FtsZ (41). Low GTPase activity indicates two possibilities: weak hydrolysis of FtsZ or weak interactions with FtsZ. FtsZ, which has a weak hydrolytic ability, remains bound to GTP and is less likely to depolymerize. In contrast, FtsZ, which weakly interacts with GTP, was less polymerized.

### Structures of FtsZ filaments

Previous electron microscopy observations have shown that FtsZ filaments of *E. coli* and *B. subtilis* are formed at 4–5 nm wide (42, 43). The width of the SeFtsZ filaments (5.7 ± 1.2 nm) we measured was similar in value. In addition, the filament length was shorter than that of *E. coli* and *B. subtilis*. These short filaments indicate that SeFtsZ does not easily interact with GTP. The SeFtsZ filaments formed ring structures. The ring structures with a diameter of ∼300 nm were also observed in *B. subtilis* FtsZ (44). The ratio of the FtsZ ring size in *Spiroplasma* to that in *B. subtilis* was the same as the cell width. This suggests that the curvature of FtsZ filaments depend on cell morphology.

### Effects of SepF for FtsZ polymerization in *Spiroplasma*

A previous study reported that *B. subtilis* SepF reduced the GTPase activity of FtsZ by stabilizing FtsZ filaments (14). It is believed that the stabilization of FtsZ filaments inhibits FtsZ depolymerization, and the reduction of the FtsZ monomer prevents new polymerization. In this study, SeSepF promoted the polymerization of SeFtsZ but did not affect its GTPase activity, suggesting that SeSepF did not reduce SeFtsZ depolymerization. In *B. subtilis*, the SepF ring controls the alignment and bundling of FtsZ filaments (15, 16, 18). The predicted structure of the SeSepF dimer in AlphaFold2 was different from that of the SepF dimer in *B. subtilis* (Fig. 7), and the ring-like structure of SeSepF was not observed by EM. We showed that SeFtsZ monomers interacted more strongly with SeSepF than the polymers. The critical concentration required for FtsZ filament formation is higher in *E. coli* than in *Spiroplasma melliferum* (41). Due to the high critical concentration, several SeFtsZ monomers were present around the polymerized SeFtsZ. Since SeSepF easily interacts with the SeFtsZ monomer, the release of SeSepF from the SeFtsZ filaments is facilitated, leading to the formation of unstable SeFtsZ filaments.

**Figure 7.**
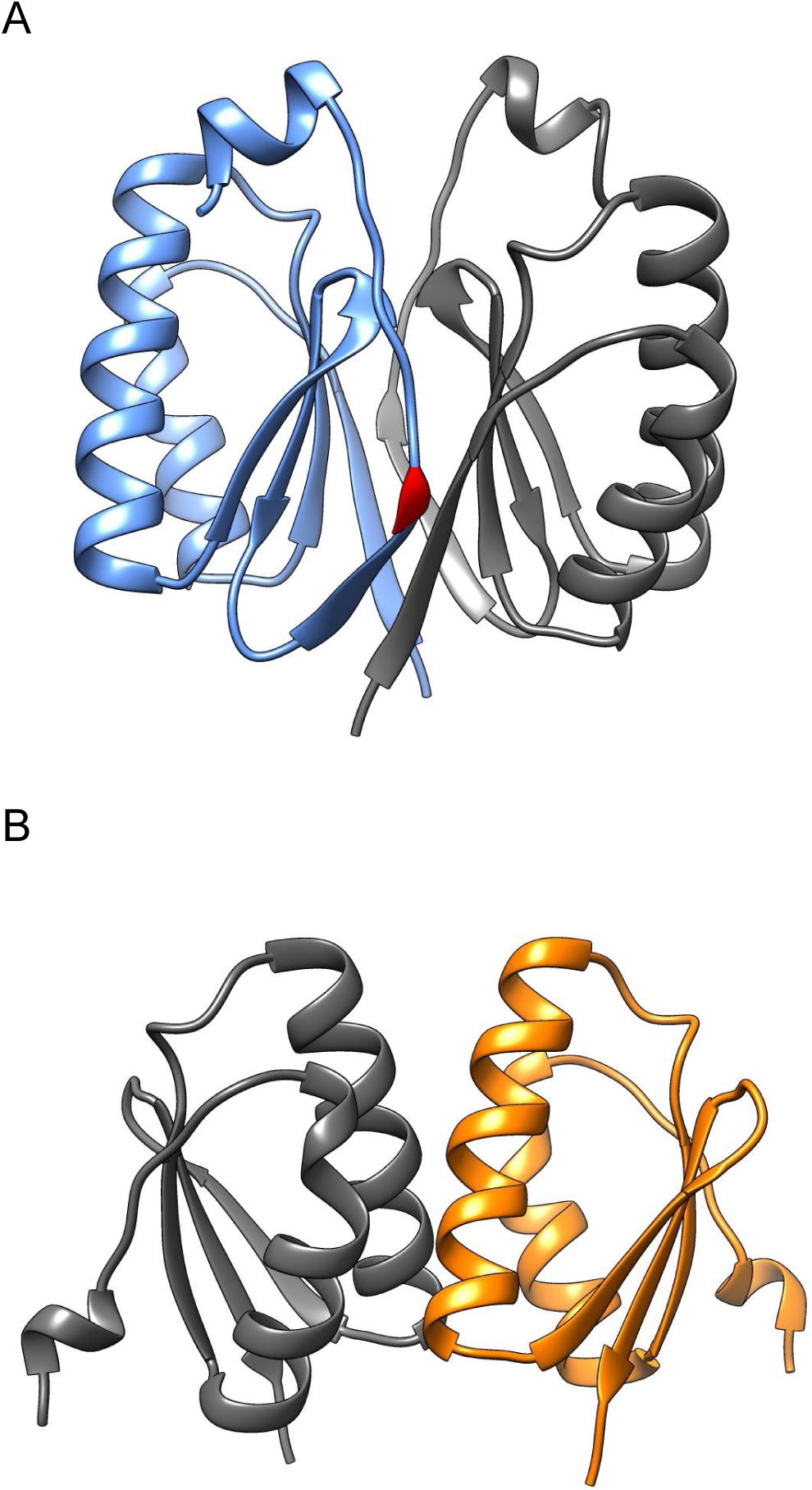
Predicted 3D structure model of SepF dimer. (A) The G137 (shown in red) is important for interaction of *B. subtilis* SepF. (B) SeSepF is shorter than *B. subtilis* SepF and lost G137 residue.

### Cell division of Spiroplasma eriocheiris

A previous study reported that Mollicutes FtsZ produced in *E. coli* localizes to the cell pole and inhibits cell division (45, 46). Here, we report the movement of SeFtsZ in *E. coli* L-form cells, in which *E. coli* FtsZ does not function. SeFtsZ formed ring- and tube-like structures in L-form cells but did not constrict the cell membrane. Next, we showed that when SeFtsZ and SeSepF were coexpressed in L-form cells, SeFtsZ localized to the cell membrane and formed ring-like structures. Ring-like structures constricted cell membranes. The SeFtsZ ring was similar to that of *E. coli* FtsZ when expressed in liposomes (28, 47, 48). Our findings also showed that SeFtsZ formed Z rings in a SepF-dependent manner in *E. coli* cells. To our knowledge, this is the first report of Z-rings derived from Mollicutes. However, the SeFtsZ ring may require a physical force to separate the cell membrane. The motility of *Mycoplasma* is involved in the cell division process (49, 50). *S. eriocheiris* also divides using physical forces generated by swimming motility, and SeFtsZ determines the division sites by narrowing a part of the cell.

## Experimental Procedures

### Bacterial strains and plasmid constructions

*E. coli* MG1655 is the wild-type strain (51). *E. coli* Rosetta (DE3) cells were used for protein production. *Spiroplasma* genomic DNA was purified using the Wizard Genomic DNA Purification Kit (Promega, Madison, WI, USA). The *ftsZ* and *sepF* genes were amplified using genomic *Spiroplasma* DNA as a template. The TGA codon in *SeftsZ* and *SesepF* was changed to a TGG codon for expression in *E. coli*. *SeftsZ* and *SeSepF* were inserted into the pET28a vector at the NdeI and EcoRI sites, yielding pSP2 and pSP3, respectively. SeFtsZ and SeSepF mutants were generated by site-directed mutagenesis using polymerase chain reaction (PCR). *SeftsZ^L272E^*, *SeSepF^G116R^*, *SeSepF^G123S^* were inserted into the pET28a vector using the NdeI and EcoRI sites, yielding pSP71, pSP68, and pSP69, respectively. SeFtsZ and SeSepF mutants were created by replacing pSP71, and pSP68, pSP69 with pSP27. *SeftsZ^L272E^*, *SeSepF^G116R^*, *SeSepF^G123S^* were inserted at the EcoRI and PstI sites of pBAD33-MCS3 to yield pSP89, pSP90, and pSP91, respectively. pBAD24 and pBAD33 were excised using ClaI-HindIII (52). The small fragment of pBAD24 was replaced with the corresponding fragment of pBAD33, and the resulting plasmid was named pBAD33-MCS3 (Yamaguchi, unpublished). The PstI-ScaI fragment of pDSW208 was replaced with the corresponding fragment of pDSW207 (53), yielding pRU1565. msfGFP was amplified using RU1514 (mreB-msfGFP^SW^) as a template, and the PCR product was designed to carry EcoRI and SacI sites at its 5’ and 3’ ends. The PCR products were digested with EcoRI and SacI and inserted into the corresponding site of pRU1565 to yield pRU1563. msfGFP was amplified using pRU1563 as the template. msfGFP fragment that contains the overlap sequence at the 3’ end fused to SeftsZ by PCR. The *gfp-SeftsZ* and *gfp-SeftsZ-SesepF* were inserted into the EcoRI and PstI sites of pBAD33-MCS3 to yield pSP23 and pSP27, respectively.

### Protein purification

Rosetta (DE3) strains carrying a plasmid encoding SeFtsZ, SeSepF, or their mutants were grown in an LB medium containing Kanamycin at 37 °C until the OD_600_ reached 0.6. Then, 100 μM isopropyl-β-D-1-thiogalactopyranoside (IPTG) was added to the culture to induce expression of *SeftsZ* or *SesepF,* and cells were incubated for 13 h at 25 °C. The cells were collected by centrifugation at 7,000 × *g* at 4°C. The pellet was resuspended in a binding buffer (20 mM Tris-HCl pH 7.4, 500 mM NaCl) and sonicated. The lysate was centrifuged at 22,000 × *g* for 20 min at 4 °C. The supernatants were passed through a 0.22-μm filter. The sample solutions were loaded onto a HisTrap HP column in an ÄKTA pure 25 (Cytiva, Tokyo, Japan). The column was washed with binding buffer and eluted with a linear gradient of 10–500 mM imidazole in the binding buffer. The eluted fractions were separated using SDS-PAGE (12%), followed by CBB staining. His tags were removed using a Thrombin Cleavage Capture Kit (Merck, Darmstadt, Germany).

### Sedimentation assay

The purified SeFtsZ was mixed with or without SeSepF in a polymerization buffer (50 mM Tris-HCl pH 7.4, 100 mM KCl, 10 mM MgCl_2_, 2 mM GTP or GDP) and incubated for 30 min at 25 °C. The final concentration of SeFtsZ and SeSepF were 12 μM. The polymerized SeFtsZ was centrifuged at 4 °C for 15 min at 350,000 × *g*. The pellet and supernatant fractions were analyzed using SDS-PAGE, followed by CBB staining.

### Bio-layer interferometry

Protein–protein interactions were measured using the Octet N1 system (Sartorius, Göttingen, Germany). His-SeFtsZ and His-SeSepF were diluted in BLI buffer (20 mM Tris-HCl pH 7.4, 100 mM KCl, 10 mM MgCl_2_, 0.02% Tween-20). At the loading step, 10 μg/mL His-SeFtsZ and His-SeSepF were immobilized to the Ni-NTA biosensor tips. The sensor tip was dipped into different concentrations of SeFtsZ (0–100 μg/mL) for the association step and moved into the BLI buffer for the dissociation step. The data were analyzed using Octet N1 software.

### Electron microscopy

Polymerized SeFtsZ was placed on carbon-coated glow-discharged grids for 5 min at room temperature. After removing the solution, grids were stained with 2% uranyl acetate. The images were acquired using a JEM1010 EM (JEOL, Tokyo, Japan) equipped with a FastScan-F214(T) charge-coupled device camera (TVIPS; Gauting, Germany).

### GTP hydrolysis assay

The GTPase activity of SeFtsZ was measured using malachite green assay. Purified SeFtsZ was incubated with 1 mM ATP at 25 °C. GTP hydrolysis of SeFtsZ was terminated at various reaction times, and inorganic phosphate (Pi) levels were detected using malachite green. The absorbance was measured at 620 nm using a microplate reader. Phosphate concentrations were determined using a phosphate standard curve.

### Microscopic observation

MG1655 cells producing GFP-SeFtsZ with or without SeSepF were grown to log phase in L medium. Cells were observed using an Axio Observer (Zeiss), and sectioning images were captured along the z-axis at 0.27 mm intervals and treated using a deconvolution algorithm. MG1655 cells producing GFP-SeFtsZ with or without SeSepF were grown in NB/MSM medium (0.1% Lab-Lemco powder, 0.2% Yeast extract, 0.5% Peptone, 0.5% NaCl, 40 mM MgCl_2_, 1 M sucrose, 40 mM maleic acid, pH 7.0) at 30 °C until OD_600_ reached 0.4. The cell suspension was loaded onto a CellASIC ONIX B04A microfluidic plate (Merck). The air and liquid in the chamber were replaced with nitrogen gas and NB/MSM medium supplemented with 300 μg/mL penicillin G, respectively. Microfluidic plates were observed using a phase-contrast microscope (Zeiss, Oberkochen, Germany). The microscopic images were analyzed using Fiji (NIH, Bethesda, MD, USA) or Zen (Zeiss).

## Data availability

The data supporting the conclusions of this article will be made available by the authors.

## Acknowledgments

We thank all members of the Shiomi Lab for their helpful discussions and suggestions.

## Abbreviations and nomenclature

PBPs: penicillin-binding proteins
CBB: Coomassie brilliant blue
Kd: dissociation constant
EM: electron microscope
msfGFP: monomeric superfolder GFP
Pi: inorganic phosphate

